# Repeat-induced point mutation and gene conversion coinciding with heterochromatin shape the genome of a plant pathogenic fungus

**DOI:** 10.1101/2022.11.30.518637

**Authors:** Jovan Komluski, Michael Habig, Eva H. Stukenbrock

## Abstract

Meiosis is associated with genetic changes in the genome - via recombination, gene conversion, and mutations. The occurrence of gene conversion and mutations during meiosis may further be influenced by the chromatin conformation, in analogy to what is known for mutations during mitosis. To date, however, the exact distribution and type of meiosis-associated changes and the role of the chromatin conformation in this context is largely unexplored. Here, we determine recombination, gene conversion, and *de novo* mutations using whole-genome sequencing of all meiotic products of 23 individual meioses in *Zymoseptoria tritici*, an important pathogen of wheat. We could confirm a high genome-wide recombination rate of 65 cM/Mb and see higher recombination rates on the accessory compared to core chromosomes. A substantial fraction of 0.16% of all polymorphic markers was affected by gene conversions, showing a weak GC-bias, and occurring at higher frequency in regions of constitutive heterochromatin, indicated by the histone modification H3K9me3. The *de novo* mutation rate associated with meiosis was approx. three orders of magnitude higher than the corresponding mitotic mutation rate. Importantly, repeat-induced point mutation (RIP), a fungal defense mechanism against duplicated sequences, is active in *Z. tritici* and responsible for the majority of these *de novo* meiotic mutations. Our results indicate that the genetic changes associated with meiosis are a major source of variability in the genome of an important plant pathogen and shape its evolutionary trajectory.

**Importance:** The impact of meiosis on the genome composition via gene conversion and mutations is mostly poorly understood, in particular for non-model species. Here, we sequenced all four meiotic products for 23 individual meioses and determined the genetic changes caused by meiosis for the important fungal wheat pathogen *Zymoseptoria tritici*. We found a high rate of gene conversions and an effect of the chromatin conformation on gene conversion rates. Higher conversion rates were found in regions enriched with the H3K9me3 – a mark for constitutive heterochromatin. Most importantly, meiosis was associated with a much higher frequency of *de novo* mutations than mitosis. 78% of the meiotic mutations were caused by repeat-induced point mutations – a fungal defense mechanism against duplicated sequences. In conclusion, the genetic changes associated with meiosis are therefore a major factor shaping the genome of this fungal pathogen.

## Introduction

Meiosis is an important mechanism of genome evolution as it generates genetic variability for selection to act upon. Since all changes to the genome that occur prior to and during meiosis are potentially affecting the germline, meiosis is a pivotal mechanism in shaping the evolutionary trajectory of sexually propagating species. Three major classes of genetic changes are associated with meiosis; i) recombination, ii) gene conversion and iii) meiotic mutations. Recombination is the reciprocal exchange of information between homologous chromosomes during meiosis (1). Canonical meiosis is initiated by the formation of double-strand breaks (DSBs) by the topoisomerase-like protein Spo11 at many genomic locations (2, 3). Indeed, these DSBs, which are resected to generate single-stranded DNA overhangs that can invade the homologous chromosome, are thought to guide chromosome pairing in many species (3, 4). Most DSBs are repaired and resolved as non-crossover (NCO) events which in some occasions are associated with gene conversions, while few DSBs will be resolved to crossover (CO) events and the reciprocal exchange of larger chromosome portions between homologous chromosomes (4–8). Interestingly, recombination rates, i.e. the rate of reciprocal exchanges of chromosome sections during meiosis, vary considerably between species (9, 10). The highest CO frequencies are found in fission yeast where an average of 11-19 COs per chromosome pair by far exceed the minimum of one crossover event per chromosome considered to be required for proper chromosome pairing and segregation in most species (11). Recombination rates also vary along chromosomes, with crossovers occurring in hotspots (12, 13) and being mostly absent in centromeric regions (14, 15). Where crossovers are occurring seems to be affected by the synaptonemal complex, a protein structure that forms along meiotic chromosomes, as well as the chromatin structure that appears to influence the location of the DSBs (9). Accessible chromatin appears to be a hotspot for DSBs, as shown in *Saccharomyces cerevisiae* where DSBs are primarily located within regions of accessible chromatin generally found at gene promotors (16). Generally, heterochromatic marks are associated with lower recombination rates, while euchromatic marks are associated with elevated recombination rates (3, 9, 17–20). During the last decade, the use of population data for the determination of recombination rates became feasible based on the rapidly increasing availability of whole genome sequencing data (21–24). However, progeny analysis and tetrad analysis are still required to analyze all the genetic processes associated with crossover events.

Gene conversion is also one of the possible outcomes of DSB formation and resolution during meiosis, but, in contrast to recombination, gene conversion directly affects the allele frequency. Gene conversion describes the non-reciprocal (i.e. unidirectional) transfer of a sequence from one locus (the donor) to a different genetic locus (the acceptor) (25). Gene conversions can either be interallelic or non-allelic (also called interlocus). The first will result in changes in the allele frequency while the latter is (next to unequal crossovers) involved in gene duplication, gene expansion, and homogenization of gene families and has for example been observed in gene families involved in host-pathogen interactions (25–28). Gene conversion is initiated by DSBs, followed by resection of the DSB end and the invasion of the single-stranded tail into homologous sequences. Sequence differences between the two homologous sequences will result in partially mismatched heteroduplex DNA (5, 25, 29). If the mismatch is repaired using the information of the invading DNA the acceptor allele will be changed to the donor allele and hence result in gene conversion as manifested by a 3:1 rather than 2:2 segregation pattern in the resulting products of a single meiosis – a tetrad. Heteroduplex DNA and repair also occur during crossover (CO) events and hence gene conversions can be categorized into those associated with CO (CO-GC) and those associated with non-crossover NCO (NCO-GC) (29). Rates of gene conversion vary considerably between species (30, 31) which appears to be mainly influenced by the tract length (i.e. the length of the sequence containing converted markers) and recombination rates (5, 31, 32). Gene conversion in some species seems to be GC-biased, probably caused by the GC-biased repair of A:C and G:T in the heteroduplex DNA (33, 34) which is assumed to have important consequences on the equilibrium GC content of genomes (31, 35). Biased gene conversion may however not be universally important as it was found not to occur in some fungi as well as in some plant and algae species (21). Although chromatin configuration and hence the histone modifications are assumed to affect the rate of gene conversions, such an association was so far not identified.

Finally, meiosis is also associated with mutations that occur before or during meiosis. Since mutations on average are considered to be deleterious, mutation rates, in general, are low but can also differ greatly between species (36). Meiosis-associated mutation rates, in turn, can differ greatly from the corresponding mitotic mutation rate in the same species indicating different mechanisms and/or constraints. For example, the germline mutation rate in humans and mice is 1.2 × 10^−8^ or 5.7 × 10^−9^ per nucleotide per generation, respectively, two orders of magnitude lower than the corresponding mitotic mutation rate (10, 37, 38). Here, germline mutations might be linked to DSBs and their repair (39) with a higher number of mutations occurring in the vicinity of recombination events (40). Estimates of meiosis-associated mutation rates in different fungal species also vary considerably. In *S. cerevisiae*, the meiosis-associated mutation rate is 8 × 10^−8^ per bp per cell generation (39), much higher than the mitotic mutation rate of 3.3 × 10^−10^ per bp per cell division (41). In *Neurospora crassa*, the meiosis-associated mutation rate is very high at 3.38 × 10^−6^ per bp per generation (42) contrasting with a much lower mitotic mutation rate of 6.7 × 10^−10^ per bp per cell division (43). Interestingly, this extremely high meiotic mutation rate in *N. crassa* is caused by repeat-induced point mutation (RIP), a fungal defense mechanism against duplicated sequences (44, 45). RIP is restricted to haploid parental nuclei just prior to karyogamy and meiosis and acts on duplicated sequences of a minimum length of 400 bp length in *N. crassa* (46–48). Once recognized both duplicated sequences will be mutated in a C→T manner (46–48) and RIP can sometimes leak into adjacent non-repetitive regions (49, 50). RIP signatures have been detected in the genomic sequences of many fungi, however, active RIP was experimentally confirmed only in a few fungal species (42, 51). Hence, there is a growing body of evidence that suggests that the mutational processes prior to and during meiosis differ from those during mitosis. In fungi, the meiotic mutation rate appears to be higher than the mitotic mutation rate which for some fungi is assumed to be the result of RIP.

Here, we use tetrad analysis to determine genetic changes associated with meiosis in the ascomycete fungus *Zymoseptoria tritici*, a pathogen of wheat. The haploid genome of *Z. tritici* comprises 13 core chromosomes and a set of smaller accessory chromosomes (52). These non-essential accessory chromosomes carry a fitness cost (53), are enriched in the facultative heterochromatin mark H3K27me3 (54), and show a meiotic drive (55). The availability of complete tetrads for *Z. tritici* allows us here to address all three major classes of genetic changes associated with sexual reproduction. In particular, the frequency and distribution of mutations associated with sexual reproduction are unknown although *Z. tritici* has an asexual and a sexual reproductive cycle with the latter being the main source of the primary inoculum during the initial stages of the infection (56). The mitotic mutation rate in *Z. tritici* has been determined experimentally by mutation accumulation experiments at 3.2 × 10^−10^ per bp per mitotic cell division (57), which is similar in other fungi (41, 43). Although histone modifications affect the mitotic mutation rate of *Z. tritici* (57, 58) it is unknown if the distribution of meiotic recombination events, gene conversion, and meiosis-associated mutations are also influenced by these histone modifications. Finally, although the genome of *Z. tritici* shows signatures of RIP (59), RIP has so far not been demonstrated experimentally and the efficacy of this mutational mechanism in this pathogen is not known. Given the fact that 18.6 % of the genome of *Z. tritici* is represented by TEs it is plausible that RIP is less efficient in *Z. tritici* than in *N. crassa* or fails to recognize some duplicated regions (60). Here we study all major classes of meiosis-associated genetic changes in *Z. tritici* by analyzing whole genomes sequences of complete tetrads to, i) estimate recombination and gene conversion rates for core and accessory chromosomes; ii) determine the association between recombination and gene conversions with chromatin modifications; and iii) estimate meiotic mutation rates in *Z. tritici*. The use of tetrads allowed us to detect and describe the effects of active RIP and generate a fine-scale map of recombination and gene conversion events and its association with chromatin modifications.

## Results

### Accessory chromosomes show higher recombination rates

To determine the distribution of recombination events during meiosis in *Z. tritici*, we used previously published tetrads and obtained whole genome sequences for the tetrad progenies (61). To this end, we included 23 tetrads comprising four ascospore isolates, totaling 92 genomes. An average of **1**8772 SNPs (0.3% of all analyzed genomic sites) per tetrad was used for the analysis (see Materials and Methods). From this data, we identified individual recombination events with the CrossOver tool from the Recombine package (62) and calculated the recombination rate. A total of 1138 crossover events were observed, resulting in a genome-wide recombination rate of 65 cM/Mb consistent with previously published estimates of the recombination rate in this fungus. Intriguingly, the recombination rate was significantly higher on accessory chromosomes (92.7 cM/Mb) than on the core chromosomes (62.6 cM/Mb) (Fig 1A, Fig S1A, Table S1A-B). The recombination rate was negatively correlated with the chromosome length (Pearson’s R=-0.76, p=0.00017) (Fig 1B) but the absolute number of crossovers per chromosome was positively correlated with the chromosome length (Pearson’s R=0.96, p=6.2 × 10^−11^) (Fig S1B). When correlating the recombination rates with regions enriched in heterochromatin marks we found that regions enriched in heterochromatin marks (H3K27me3 or H3K9me3) were associated with higher recombination rates whereas the euchromatin marks H3K4me2 did not show a higher recombination rate than regions lacking all three marks (Fig S1C).

**Fig 1.**
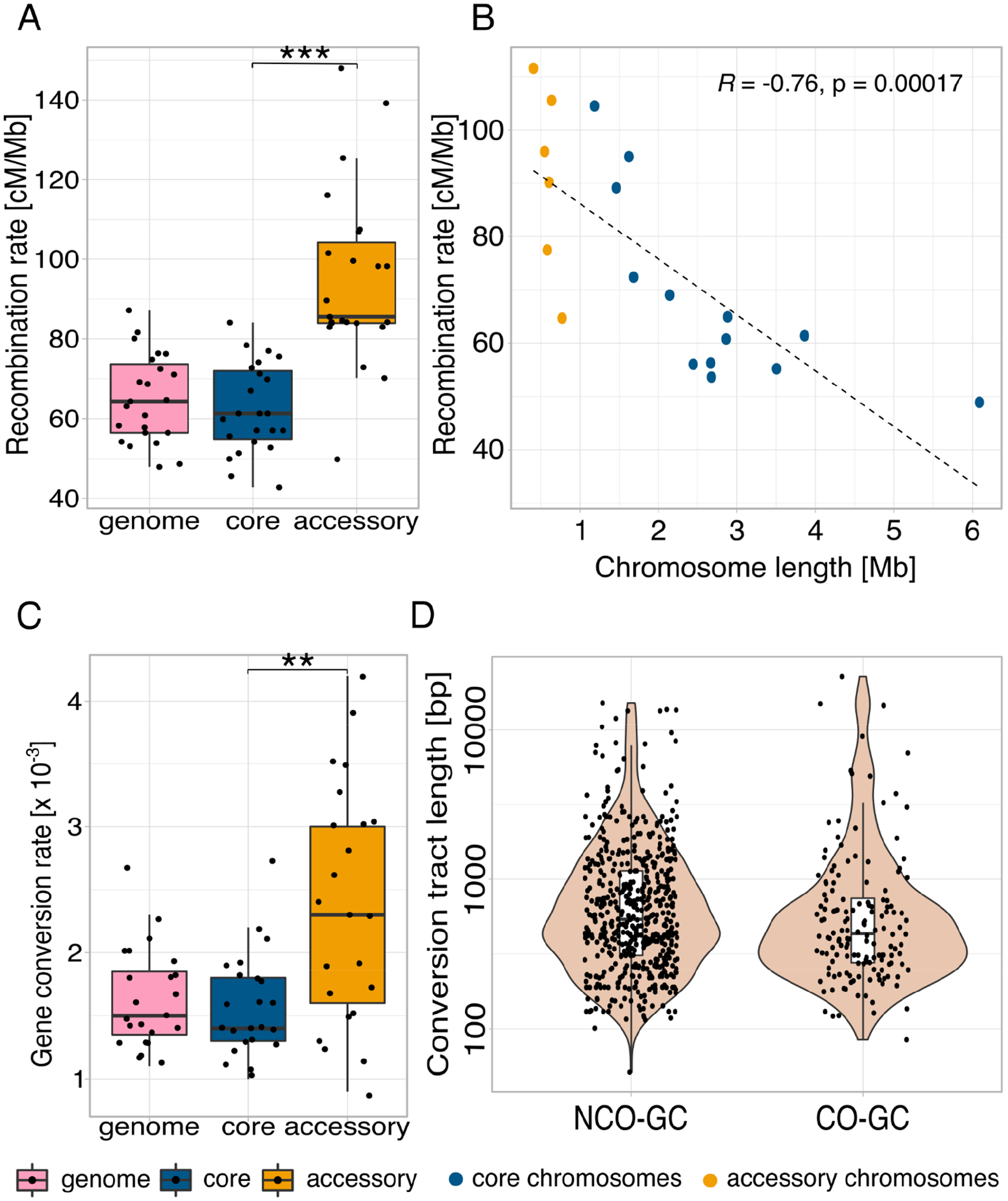
Comparison of gene conversion rates and recombination rates for different genome compartments for 23 complete tetrads. **A** Recombination rates for the genome and on core and accessory chromosomes. **B** Correlation between recombination rate and chromosome length. **C** Gene conversion rates for the genome and core and accessory chromosomes. **D** Violin plot of non-crossover associated gene conversions (NCO-GCs) and crossover associated gene conversions (CO-GCs) tract lengths detected in the study. The tract lengths of the gene conversion events spanning TEs are not shown. **A, C** *p*-values of paired Wilcoxon test are shown (\**p* < 0.05, \*\**p* < 0.005, \*\*\**p* < 0.0005).

To further assess the variation in recombination rate across the genome, we estimated the recombination rates in 20 kb non-overlapping windows. We defined hotspots as those 20 kb windows that showed a significantly (p-value<0.001) higher number of recombination events than expected by the Poisson distribution and identified a total of 52 recombination hotspots. Recombination rates per window were highly variable, ranging from 0 to 1196 cM/Mb (Fig S2). We conclude that recombination varies considerably across the genome, occurs in hotspots, and is higher on the accessory than on the core chromosomes. We considered the relevance of high recombination rates with regard to gene evolution and correlated the recombination map with the coordinates of protein-coding genes. In total, we observed 376 genes in the CO hotspots. From the 376 genes observed in the CO hotspots, six encode for effector candidates and nine genes encode CAZymes (Table S1C) which are each gene categories with a putative relevance in the pathogenicity of the fungus.

### High gene conversion rates are more uniformly distributed within the genome

The genomes of ascospore progenies resulting from one meiotic event provided us with a unique opportunity to characterize the location and distribution of gene conversion events. We, therefore, identified gene conversion events along the genome and compared the gene conversion rate for different genomic compartments. We identified a total of 890 gene conversion events comprising 712 associated with non-crossover events (Non-crossover Gene Conversion NCO-GC: 80%) and 178 associated with crossover events (Crossover Gene Conversion CO-GC: 20%) (Table S2A). We distinguished the gene conversion events on core and accessory chromosomes of *Z. tritici* and found that 755 gene conversion events were located on core and 135 on accessory chromosomes (Table S2B). We next explored the general patterns of converted SNPs. We observed a weak GC-bias in the gene conversions (binomial test, p=0.0215, Fig 2A). Based on genome-wide SNPs identified among the ascospore isolates we found that the genome-wide gene conversion rate was 1.6 × 10^−3^ per SNP (Fig 1C, Table S2B). Hence, the gene conversion rate was significantly higher on the accessory chromosomes compared to the core chromosomes (paired Wilcoxon signed rank test, *p*-value=1.0 × 10^−3^) (Fig 1C, Fig 2B). Tract length is considered to be one of the main determinants of gene conversion rates, therefore, we estimated the median conversion tract length for both types of gene conversion events. The median tract length for non-crossovers (NCO-GC) was 539 bp, and 432 bp for gene conversions associated with crossovers (CO-GC), respectively (Fig 1D).

**Fig 2.**
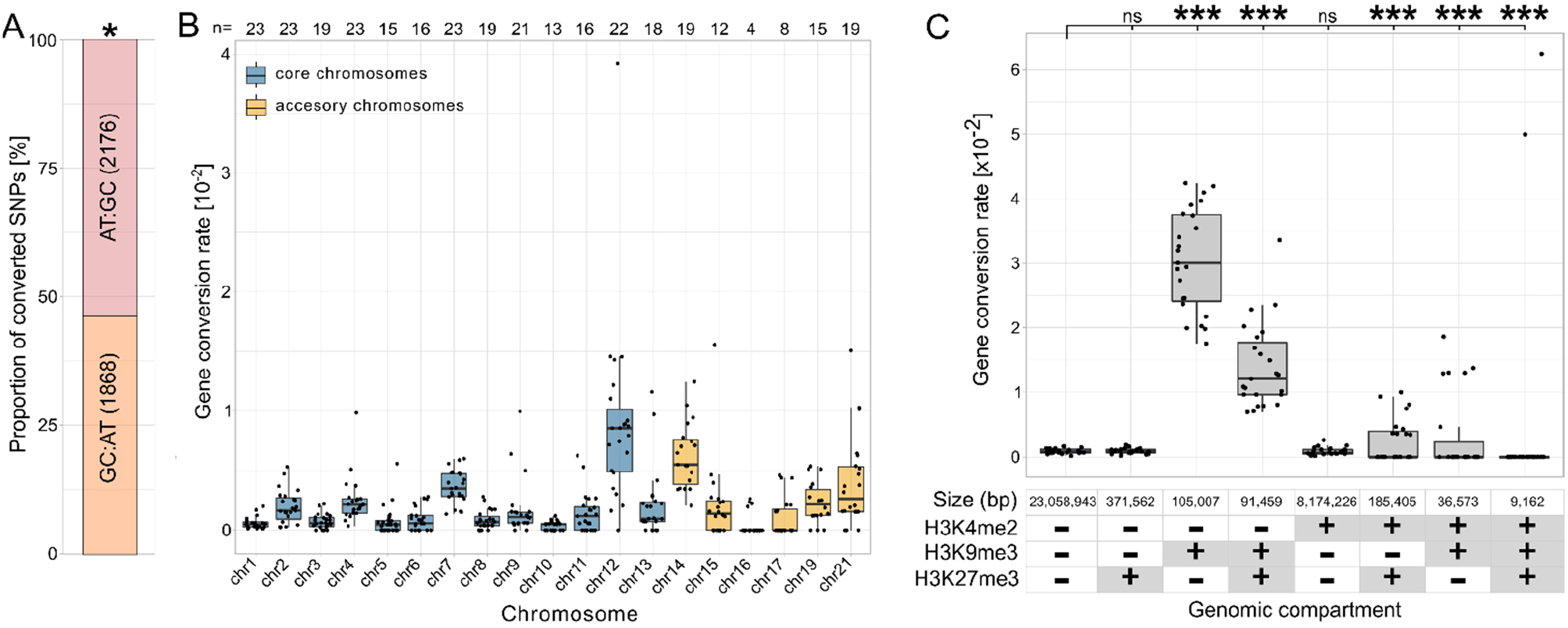
Gene conversion rate per chromosome, GC-biased gene conversion, and correlation between gene conversion rates and histone modifications. **A** GC-biased gene conversion in *Z. tritici*. The stacked barplot shows the proportion of AT to GC converted markers and the proportion of GC to AT converted markers. Binomial test *p-*values are shown (\**p* < 0.05, \*\**p* < 0.005, \*\*\**p* < 0.0005). **B** Gene conversion rate per chromosome is calculated as the proportion of converted markers from the total number of markers on the respective chromosome. The numbers above box plots show the number of tetrads with gene conversion detected on the respective chromosome. Box plots display center line, median; box limits, upper and lower quartiles; whiskers, 1.5x interquartile range; points, rate per tetrad **C** Correlation between gene conversion rates and chromatin modifications. The gene conversion rate is calculated as the number of converted markers per region divided by the total number of markers in the region. The presence/absence of the specific chromatin modification (H3K4me2, H3K9me3, or H3K27me3, respectively) in the genomic compartment is depicted with +/-in the table below the x-axis. *χ*^2^-test *p-* values are shown (\**p* < 0.05, \*\**p* < 0.005, \*\*\**p* < 0.0005).

In the same way as we characterized the distribution of recombination hotspots along the fungal genomes, we used the map of gene conversion events to identify regions with exceptionally high gene conversion rates, here defined as gene conversion hotspots. First, to assess the distribution of gene conversion, we divided the genome into 20 kb overlapping windows and calculated the number of gene conversion events in each window, and hereby we identified 32 gene conversion hotspots with *p*-value<0.001 defined by the Poisson distribution (Fig S3). Of the 145 genes in the gene conversion hotspots two genes encode for effector candidates and seven genes encode for CAZymes (Table S2C), and we consider the increased rate of gene conversion in these putative pathogenicity-related genes as a putative mechanism of rapid adaptive evolution. Taken together, the ascospore population allowed us to precisely map and characterize gene conversion events in *Z. tritici*. We found that rates of gene conversion showed less variation across the genome than recombination rates however with both being higher on accessory chromosomes than on core chromosomes.

### Gene conversion rates are higher in regions enriched in heterochromatin modifications

Previous studies in chicken B-cell lines have identified a correlation of chromatin structure with gene conversion (63) but detailed analysis of the effect of specific histone modifications is mostly missing. Thus, to investigate the potential effect of histone modifications on the rate of gene conversion in *Z. tritici*, we conducted a correlation analyses of maps of histone modifications and gene conversion rates. We focused on three histone marks which have been previously well characterized in *Z. tritici* using chromatin immunoprecipitation of antibodies targeting specific histone modifications followed by sequencing (ChIPseq): the euchromatin mark H3K4me2, the constitutive heterochromatin mark H3K9me3 and the facultative heterochromatin mark H3K27me3 (54) (Fig 2C, Table S3). Importantly, we excluded TEs from the full-factorial analysis of gene conversions and therefore regions enriched with H3K9me3 and H3K27me3 in this analysis are not associated with TEs.

Our analyses show that higher gene conversion rates associate with regions enriched in heterochromatin marks, particularly in regions solely enriched with H3K9me3 as well as in regions enriched with both H3K9me3 and H3K27me3 (Fig 2C). The median gene conversion rate in the regions enriched with H3K9me3 was 3 × 10^−2^ per converted marker per tetrad per meiosis and the median gene conversion rate in H3K9me3 and H3K27me3 regions was 1.3 × 10^−2^ per converted marker per tetrad per meiosis. Genome regions not enriched with either of the three histone marks H3K4me2, H3K9me3, and H3K27me3 showed the lowest gene conversion rate of 0.9 × 10^−3^. In summary, the histone maps allowed us to reveal that repressive heterochromatin modifications, especially H3K9me3 are associated with higher gene conversion rates in *Z. tritici*.

### The majority of meiotic mutations are caused by RIP

Recombination and gene conversion as well as meiosis and meiosis-associated processes themselves are considered to be mutagenic. Hence, we asked to which extent *de novo* mutations had occurred in the ascospore progenies. These mutations are distinguished by being absent in both of the parental strains, but present in the ascospores of the tetrad. Meiotic mutation rates often differ from mitotic mutation rates and are associated with recombination or pre-meiotic processes. We observed a total of 526 *de novo* mutations that were absent in both of the parental strains for the ascospores of the 23 tetrads. Hence, through the sexual cycle, there were on average, 22.9 mutations per genome per generation, resulting in a mutation rate of 5.7 × 10^−7^ mutations per bp per generation (Fig 3 A, Table S4). Of these mutations that originated from the sexual cycle, 242 (42%) resided in one particular region that clearly stood out as we mapped meiotic SNP along the genome. This high number of SNPs located in a 14001 bp region on chromosome 3 (Fig 3A-C). Every one of the 242 mutations in this region on chromosome 3 was a CG:TA transition. As we further inspected this region, we noted that the 14 kb region spanning 1668370 bp to 1682371 bp on chromosome 3 showed an increased sequencing coverage corresponding to a duplication of the region relative to chromosome 3 in the parent IPO94269 (Fig 3B). This means that this duplication on chromosome 3 was present in all 23 meioses and could possibly be identified by RIP, a genome defense mechanism against such duplicated regions (45). A high number of transitions in a duplicated region is the hallmark of such active RIP. Indeed, the high number of mutations in the duplicated region and the fact that all of these mutations are CG:TA transitions indicates active RIP in *Z. tritici*. Interestingly, the duplicated region on chromosome 3 showed RIP mutations in only ten of the 23 tetrads, indicating that in the remaining 13 tetrads this duplicated region was either not detected or not acted upon by RIP (Fig S4 A).

**Fig 3.**
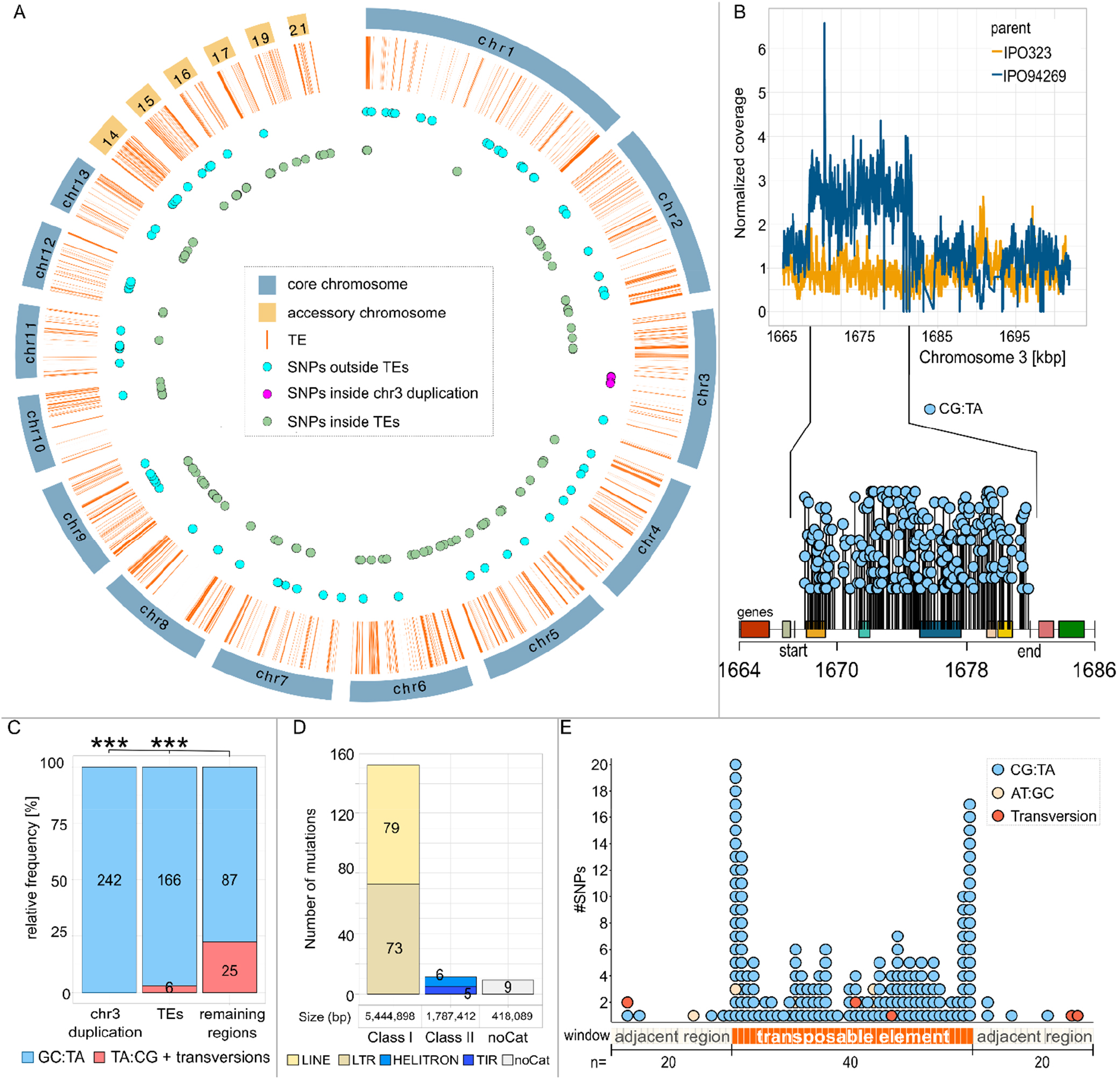
Genome-wide distribution of *de novo* mutations associated with meiosis in *Z. tritici*. **A** Circosplot of the meiotic SNPs distribution in the genomic features of *Z. tritici* (orange lines represent TEs; blue dots-meiotic SNPs outside the TEs; lilac dots meiotic SNPs inside the duplication on chromosome 3; green dots represent meiotic SNPs inside of the TEs). **B** RIP-like mutations in the duplicated 14 kb region on chromosome 3. The upper line graph shows the difference in normalized coverage between the IPO323 and the IPO94269 parent in the region on chromosome 3. The distributions of mutations in the duplicated chromosome 3 region and 10 kb upstream and downstream regions are shown in the lolliplot below the line graph. The start and the end on the x-axis of the lolliplot designate the start and the end of the duplicated region. Rectangles in different colors depict the genes located in this region. Each lollipop represents a single mutation. **C** Number of meiotic mutations in and out of the RIP active regions represented as their relative frequencies. CG:TA transitions are colored in lighter blue, and TA:CG transitions and transversions are colored in red. Fisher exact test *p-*values are shown (\**p* < 0.05, \*\**p* < 0.005, \*\*\**p* < 0.0005). **D** Number of *de novo* mutations in different classes and families of repeats (light yellow-long interspersed nuclear element (LINE); dark yellow-long terminal repeats (LTR); marine blue-HELITRON, dark blue-tandem inverted repeats (TIR); white-noCat). **E** Distribution of meiotic SNPs along TEs. Each TE was divided into 40 equal windows. Each black rectangle on the x-axis represents a window inside a TE representing 2.5% of the TE length. Beige rectangles represent windows in the regions directly adjacent to TEs. Dots above rectangles represent one mutation in each window (yellow dots-AT:GC transitions; blue dots-CG:TA transitions, red dots-transversions)

We further addressed to which extent *de novo* mutations could be the result of meiosis-associated RIP. Hereby we found that 172 of the meiotic *de novo* mutations (32% of all *de novo* mutations) were detected in transposable elements (Fig 3C), and 166 of these 172 mutations (96.5%) were CG:TA transitions; the putative signature of RIP. To examine whether these mutations were also most likely caused by RIP we determined which class of transposable elements was affected. Of the 172 *de novo* mutations in TEs 152 (88.3%) were located in Class I transposons, which replicate via an RNA intermediate and are also referred to as copy-and-paste transposons (64–66). Of these 152 mutations in Class I transposons (LINE (long interspersed nuclear elements) and LTR (long terminal repeats)) 146 were CG:TA transitions with five TA:GC transitions and one G→C transversion, whilst only eleven CG:TA transitions occurred in Class II transposable elements (Fig 3D). Class II TEs are referred to as cut-and-paste transposons, which are excised and moved to new locations in the genome and hence do not create repeated sequences (65). The mutation rate in Class I (copy-and-paste) TEs is 1.3 × 10^−6^ per bp per generation compared to 2.8 × 10^−7^ per bp per generation for Class II (cut-and-paste) TEs with the vast majority of the *de novo* mutations showing RIP-like CG:TA transitions.

Investigations in a few other ascomycete fungi have provided evidence that RIP can affect the vicinity of duplicated regions by “leakage” of mutations into these non-repeated regions (49, 50). Based on these previous studies, we here also asked if RIP mutations would be present in regions adjacent to TEs. We subdivided each TE into 40 equally sized windows (average window size = 71bp) as well as the two adjacent regions into 20 windows each (window size = 71 bp) and counted the number of *de novo* mutations in each of these windows. This approach allowed us to show that RIP mutations were not equally distributed along TE sequences. Within TEs the most distal windows showed the highest number of *de novo* mutations whereas the more central sequences were less likely to be mutated. In regions adjacent to the TEs a much lower number of mutations occurred compared to the TE sequences, but RIP-like CG:TA transitions were more frequent. It, therefore, appears that a low amount of leakage does occur in the vicinity of RIP-mutated TEs. Only 21.3% (112 mutations) of all *de novo* mutations were located in the regions outside of the transposable elements and the 14 kb region on chromosome 3. These *de novo* mutations in non-TE and non-duplicated regions comprised a significantly higher proportion of TA:CG transitions and transversions (22.3%) (Fig 4E).

The segregation pattern of the *de novo* mutations varied between the different compartments with those in TEs showing a higher proportion of 1:3 segregation (Fig S4 B) than those in the 14kb duplication or the mutations outside TEs. The segregation pattern of *de novo* mutations can indicate at what stage a DNA lesion or mismatch may have occurred and at what stage this lesion or mismatch (possibly caused by RIP) was resolved. A 2:2 segregation indicates that the mismatch was resolved prior to the replication cycle of the meiosis, while a 1:3 segregation indicates that the mismatch was resolved only during replication or later. Hence the higher proportion of 1:3 segregation pattern observed in the TEs could indicate a delayed resolution of the mismatches introduced by RIP in the heterochromatic TEs compared to the resolution in other genomic compartments. In summary, RIP has been proposed to be an important player in the genome evolution of *Z. tritici*. However, this is the first experimental evidence for active RIP in this fungus and we observed that the large duplication was not always mutated by the RIP mechanism and that the total number of mutations introduced by RIP was low - leaving many cytosines in the duplicated regions unaffected - which together could indicate a low efficiency of RIP in *Z. tritici*.

## Discussion

Here we used tetrad analysis to estimate recombination rates, gene conversion rates, and *de novo* mutation rates associated with meiosis from 23 individual meioses in *Z. tritici*. The ability to dissect genetic events in individual ascospore progenies isolated from tetrads provided us with highly precise maps of meiosis-associated changes along the fungal genome. Our results show i) higher gene conversion rates and recombination rates on accessory chromosomes compared to core chromosomes, ii) a correlation of recombination rates and gene conversion rates with histone marks associated with heterochromatin, and iii) elevated *de novo* mutation rates during sexual reproduction caused by active RIP and RIP-mutations leaking into regions adjacent to duplications.

Our dissection of recombination events during individual meiotic events allows us to confirm the previously reported high recombination rates in *Z. tritici* - on average 65 cM/Mb. Interestingly, 35 of the 52 recombination hotspots detected in this study overlapped with previously reported recombination hotspots in *Z. tritici*, which used distinct isolates from Switzerland, indicating that recombination hot spots may be determined by certain domains or conserved marks along the genome (22). In contrast to previous studies based on population genomic data (21), we show that the rate of recombination in fact is higher on accessory chromosomes compared to core chromosomes (92.7 cM/Mb and 62.6 cM/Mb, respectively) and negatively correlated with chromosome size. Hence, our results are in line with the general observation that smaller chromosomes tend to have higher recombination rates (67). Earlier population-based recombination rate studies in *Z. tritici* showed, however, lower recombination rates as computed as *rho* for accessory chromosomes (*rho* = 0.001) than on core chromosomes (*rho*=0.024) (21). *Rho* is the product of the actual recombination rate and the effective population size Ne, and we speculate that the discrepancy in recombination rate measures may reflect the lower effective population size of the accessory chromosomes compared to the core chromosomes (21). Intriguingly, we could not identify any crossovers on an accessory chromosome in nine instances, despite the presence of the respective homologous chromosomes. A minimum of one crossover per homologous chromosome pair is considered to be required for proper segregation of the homologous chromosomes (4, 68). Indeed, in four of these nine instances, we observed such segregation errors. However, in the remaining five instances the accessory chromosomes properly segregated despite the absence of crossovers. Hence, we conclude that crossovers are not essential for proper chromosome segregation in *Z. tritici. spo11* deletion mutants in *S. cerevisiae* and *Sordaria macrospora* were previously used to observe the consequences of the absence of DSBs and recombination on the segregation of chromosomes during meiosis. In these fungal species the absence of DSBs and recombination caused widespread chromosome segregation errors highlighting the importance of recombination for proper segregation (69, 70). The relatively high frequency of properly segregated, non-recombined chromosomes in *Z. tritici* indicates the presence of a non-recombination-dependent segregation system which might also be involved in the meiotic drive system of the accessory chromosomes in *Z. tritici* (61).

We predict that high levels of gene conversions in *Z. tritici* can affect allele frequencies to a higher extent than recombination and thereby shape genome composition. The genome-wide gene conversion rate of 1.6 × 10^−3^ per SNP identified in our study is approx. twenty times lower than the genome-wide gene conversion rate in *S. cerevisiae* (3.8 × 10^−2^ per SNP) (30, 31) but approx. an order of magnitude higher than the genome-wide gene conversion rate in *N. crassa* (0.7 – 2.2 × 10^−4^ per SNP) (30, 31). The variation in the gene conversion rate between species might be influenced by tract length and recombination rate since gene conversion is positively correlated with both characteristics (31, 32, 34). Our data confirm this since *Z. tritici* has shorter NCO-GC and CO-GC tract lengths and lower recombination rates (539 bp, 432 bp, and 65 cM/Mb, respectively) than *S. cerevisiae* (1681 bp, 1841 bp, and 375 cM/Mb) (31). *N. crassa* in turn shows longer NCO-GC tract lengths but shorter CO-GC tract lengths and a lower recombination rate (950 bp, 284 bp, and 20 cM/Mb, respectively) than *Z. tritici* (31). Similar to recombination, accessory chromosomes have higher gene conversion rates than the core chromosomes. The smaller size of accessory chromosomes in contrast to core chromosomes could influence the gene conversion rate on accessory chromosomes since smaller chromosomes tend to have higher rates of gene conversion (31).

We identified a strong association between chromatin modifications and recombination and gene conversion rates. We speculate that this could be caused by two non-exclusive mechanisms: The impact of histone modifications on the location of DSBs and or its effect on DSB and heteroduplex repair. Although a direct correlation between DSB formation and H3K4me3 has been shown in *S. cerevisiae* (16) the underlying causality of this correlation and to which extent the correlation also applies to other organisms is unclear (71, 72). Local chromatin conformation can influence how DSBs and heteroduplex DNA are repaired and thereby influence the later stages leading to recombination and gene conversion (73–75). In contrast to *S. cerevisiae* we see an association of heterochromatin as marked with H3K9me3 and H3K27me3 with recombination in *Z. tritici*. Similarly, and again in contrast to the scenario in yeast, we observe that gene conversion rates are highest in heterochromatin regions in *Z. tritici*. This indicates that chromatin conformation might be affecting gene conversion primarily via its effect on DSB and heteroduplex repair. DSBs can be resolved by synthesis-dependent strand annealing (SDSA) resulting in a homologous recombination-mediated pathway and therefore NCO events (76, 77). Our results show elevated gene conversion rates in regions enriched in heterochromatin marks, primarily in H3K9me3. Several studies imply that H3K9 di-or trimethylated (H3K9me2/3) heterochromatin is promoting homologous recombination (HR) (77–81). The formation of the DNA double-strand promotes the stabilization of the chromatin structure by H3K9me3 and activates DSB-signaling proteins (82). The role of H3K9me3 in promoting homologous recombination could imply that the correlation between H3K9me3 and the gene conversion rate that we observed in our study is caused by the effect of H3K9me3 on recombination. However, 80% of the gene conversion events that we detected are associated with non-crossover events (NCO-GC). This indicates that the correlation of H3K9me3 and gene conversions is not caused by recombination but possibly by the effect of histone modifications on the repair of DSBs and heteroduplex DNA. In contrast to DSBs in regions enriched with H3K9me3 regions, DSBs in regions enriched with H3K27me3 are found to be frequently repaired by microhomology-mediated end joining (MMEJ), a non-homologous repair pathway that does not promote homologous recombination and therefore does not lead to gene conversion or crossover (83, 84). We find support for a similar effect of chromatin modifications on DSB repair in our study. Indeed, increased repair of DSBs via a non-homologous pathway in the H3K27me3-enriched regions could explain the reduction in the gene conversion rate in the regions enriched in both H3K9me3 and H3K27me3 regions (1.3 × 10^−3^ per SNP) in comparison to regions solely enriched in H3K9me3 (2.8 × 10^−3^ per SNP) observed here. In conclusion, we see indications that histone modifications could affect gene conversion rates mainly via their effect on the DSB and heteroduplex repair.

*De novo* mutations associated with meiosis occured at a rate of 5.7 × 10^−7^ per bp per generation in *Z. tritici* which is approximately three orders of magnitude higher than the mitotic mutation rate (3.2 × 10^−10^ per site per cell division) which we previously determined in a mutation accumulation experiment (57). Similarly, higher meiotic than mitotic mutation rates have been reported in other fungi like *S. cerevisiae* and *N. crassa* (39, 41–43). In *N. crassa*, the difference between mutation rates during mitosis and meiosis is mostly due to RIP, a fungal defense mechanism against duplicated DNA sequences that occurs in the haploid nuclei just prior to meiosis and that induces CG:TA transitions in duplicated sequences and transposable elements (42, 46). In *Z. tritici* signatures of past RIP have been found by analyses of genome data (59, 85), and here we can confirm that RIP is an active mechanism in *Z. tritici*. This is very evident as 77% of the *de novo* mutations associated with meiosis located in Class I transposable elements (copy-and-paste elements) and in the 14001 bp long region on chromosome 3 that is duplicated in the IPO94269 parent. Interestingly, RIP in *Z. tritici* is not consistently efficient in mutating duplicated sequences as we find that duplicated sequences in some of the tetrads were not mutated. Until now, RIP was experimentally demonstrated in only a few fungal species (86– 91), with lower efficiency in *Leptosphaeria maculans* and *Podospora anserine* compared to the highly efficient RIP in *N. crassa* (88, 92). We consider that variation RIP efficiency may be a more common phenomenon that reflects a trade-off between the evolutionary costs of the mutations introduced by RIP and the evolutionary costs of TEs.

We find that Class I (copy-and-paste) TEs show a much higher RIP mutation rate than Class II (cut-and-paste). This observation is expected as the transposition mode of the Class I TEs results in duplicated sequences that are recognized by the RIP machinery (64). Interestingly we find that the RIP mutations are not equally distributed along the TE sequences. The most distal windows comprising 5% of the length of the TEs comprise 35% of all *de novo* meiotic mutations that occurred within the TEs. This high frequency of *de novo* mutations at the ends of TEs for LINE transposons could be due to the mode of transposition of Class I transposons. For these transposons the reverse transcription of the RNA intermediate and the integration of the resulting DNA starts from the 3’end -usually the polyadenylated tail (93). This process of reverse transcription and integration is frequently found to be disrupted which results in only the terminal fragments of the LINE transposon becoming integrated into the target site (93) and thereby the terminal parts of LINE transposons are more likely duplicated. LTRs transposons contain terminal repeats of 200 bp to 500 bp in length and therefore already contain duplicated sequences in close vicinity (66). These duplicated terminal sequences in Class I transposons seem to be prominent targets for RIP. As shown in other studies, and notably in *N. crassa*, the mutagenic effects of RIP leaks into adjacent regions from the terminal sequences of TEs (46). Leakage of RIP from duplicated regions into adjacent regions has been proposed to play a role in rapid adaptive evolution of effector genes involved in host-pathogen interactions (49, 94, 95). Indeed, in the pathogen *L. maculans*, effector genes locate in the vicinity of TEs and show signs of rapid evolution due to RIP mutations and particular position of these genes. Taken together, we show that RIP not only affects transposable elements in *Z. tritici* but also leaks into adjacent regions which, by the nature of the very compact genome of this fungus, can affect the mutational environment of closely located genes.

In conclusion, we show that gene conversion is correlated with histone modifications in *Z. tritici* and that RIP is active, albeit at a lower efficiency than in *N. crassa*, in this fungus affecting duplicated sequences as well as TEs and leaking into adjacent regions. As a result, meiotic mutation rates for *Z. tritici* are three orders of magnitude higher than the mitotic rates demonstrating the major impact that genetic changes associated with meiosis have on the genome composition of this important plant pathogen.

## Materials and Methods

### Fungal material

Tetrads used for the sequencing analysis were obtained from the study of Habig et al. 2018 (61) and include ascospores isolated from crosses between the Dutch isolates IPO94269 and IPO323 (available from the Westerdijk Institute (Utrecht, The Netherlands) with the accession numbers CBS115943 and CBS115941) and from the crosses between IPO94269 and whole chromosome isogenic deletion strains (Δchr14-Δchr20) of the reference strain IPO323 generated in the study of Habig et al. 2017 (53). All ascospores were cultivated for DNA extraction at 18°C at 200 rpm in YMS (4 g/L yeast extract, 4 g/L malt, 4 g/L sucrose) medium for 5-7 days inoculated directly from -80°C glycerol stocks.

### Genome sequencing and data analysis

For sequencing, DNA of 84 ascospores was isolated using a phenol-chloroform extraction protocol as described previously (61). Library preparation and sequencing using an Illumina HiSeq3000 machine for the 84 ascopores were performed at the Max Planck-Genome-centre, Cologne, Germany. The Illumina read data is available in the Sequence Read Archive under the BioProject PRJNA904559. Please note that two tetrads (A03-4 and A08-1) that were sequenced in an earlier study were also included (61). The Illumina read data for these two tetrads is available at the Sequence Read Archive under the BioProject PRJNA438050. An overview of the included tetrads and ascospores and is given in Table S5. Please note that one of the originally 24 tetrads was excluded from the analysis because it showed a pattern of SNPs that were inconsistent with meiotic recombination on several chromosomes and was assumed to be not the product of a single meiosis but rather a mixture of two or more meiotic events. In addition, the parental strain IPO94269 was sequenced as described above and its Illumina reads were deposited in the BioProject PRJNA904744. The Illumina data for the IPO323 parental strains is available under the BioProject PRJNA371572.

### Mapping and SNP calling

The reads of 92 ascospores of the 23 tetrads were mapped to the IPO323 reference genome with bowtie2 (version 2.3.4.1) (96). This analysis focused only on single nucleotide polymorphisms (SNPs). To obtain a high-quality SNP dataset we performed the SNP calling with two variant callers, GATK (version 4.1.6.0) (97) and Samtools (version 1.7) (98), and SNPs with QUAL≥90 that were called by both variant callers were used for the downstream processing. Variants in regions that contain transposable elements (TEs) were removed from the analysis with bedtools intersect (version 2.26.0, option -v) (99) to avoid spurious alignments. From the remaining SNPs, only variants from regions with coverage > 5 in all four ascospores of a tetrad were used for the analysis to avoid false negatives due to the low coverage in one of the spores. Biallelic SNPs with minor allele frequency >0.9 and with QUAL>90 that were called by both variant callers were identified by overlapping variant call format (VCF) files from both haplotype callers with bedtools intersect and used as a core set of high-fidelity SNPs. VCF files of four spores from the same tetrad were merged with VCFtools (v0.1.15) with merge option (100) to create a variant file for each tetrad.

High-quality threshold and variant calling accepting only variants identified by two callers will potentially lead to false negative calls in some of the four spores and hence will affect the segregation ratio obtained for the SNPs. To reduce the risk of false negative calls we reintroduced high-fidelity SNP to any of the other ascospores of a tetrad if there was an indication that it was present but did not satisfy the quality requirements. This means that SNPs which were called in a tetrad but did not meet the quality threshold in some of the ascospores but met the quality threshold in at least one of the ascospores were re-introduced. SNPs on chromosomes that were deleted or absent in the parental strains were removed from the analysis of the respective progeny. Please see Table S5 for an overview of the number of high-fidelity SNPs included for each tetrad.

### Identification of recombination events

To detect recombination and gene conversion events in the tetrad progeny, CrossOver.py from the Recombine package (version 2.1) for tetrad analysis in yeast (62) was modified to fit the genome characteristics of *Z. tritici* (size and number of chromosomes and the location of the centromeres (54)). Input segregation files for the CrossOver program were generated from merged tetrad VCF files for each tetrad with the custom-made bash script (see Supplementary methods). Each segregation file consisted of 7 columns: the first two columns referred to the chromosome and position of the variant, the third column served as a spacer, and the last four columns referred to the presence/absence of SNPs in four spores. 0 and 1 values in the last four columns of a segregation file designated the presence or absence of a variant at a certain position compared to the reference genome. The program initially identifies COs as positions with 2:2 segregation where adjacent markers undergo a reciprocal genotype switch. Gene conversion tracts are then identified as regions of non-2:2 segregation. After the identification of recombination events, all double crossovers separated with a single SNP were filtered out. Gene conversions were filtered for tracts spanning ≥ 3 markers. The recombination rate per tetrad [cM/Mb] was calculated by the following formula:

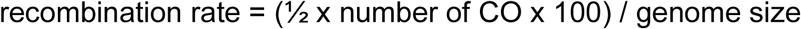

The gene conversion rate per tetrad was determined as the proportion of converted markers from the total number of markers identified per tetrad. Furthermore, tract lengths were determined with the midpoint method, i.e. the midpoint between two markers of a different class (e.g. converted vs. non-converted) was considered to be the position were the tract started or ended. Tracts spanning TEs were removed for the estimation of tract lengths and recombination rates. Recombination events and gene conversion events detected in this study are listed in the supplementary material (Table S1B-C and Table S2A-C, respectively). SNPs in TEs were disregarded for the determination of recombination and gene conversion events.

### Estimation of meiotic mutation rates

To estimate meiotic mutation rates in *Z. tritici*, genome-wide SNPs (including SNPs in TEs) satisfying the following criteria were taken into consideration: i) read depth > 5 in both parental strains and the ascospore progeny; ii) absent in both parental strains (Table S4). Before a SNP in the progeny was considered a *de novo* mutation, both parental sequencing results were manually checked to validate that this SNP was already present but not called in the parental genomes. Only SNPs in the progeny that showed no hints in the parental genomes were included in the subsequent analysis. The per bp mutation rate was calculated as the “average number of meiotic mutations per ascus” / “the reference genome size”. To verify *in silico* detected meiotic mutations, we performed Sanger sequencing of 20 randomly selected mutations from which 19 mutations were confirmed (see Supplementary Methods).

### Detection of duplications

For the detection of duplications in the parental strains, Illumina reads were quality filtered as described above and mapped onto the reference genome using SpeedSeq align followed by structural variation analysis by LUMPY (101) as implemented in the SpeedSeq package (version 0.1.2) (102). The VCF files were filtered using bcftools (version 1.6) as follows: VCF files were filtered on duplications, genotype (GT=0/1) and quality >400, and length < 50000.

## Data Availability

The Illumina read data is available in the Sequence Read Archive under the BioProjects PRJNA904559, PRJNA438050, PRJNA904744, and PRJNA371572. The *Z. tritici* IPO323 reference genome is available under the accession GCA_000219625.1.

## ACKNOWLEDGMENTS

The study was funded by a personal grant to EHS from the State of Schleswig Holstein and the Max Planck Society and in addition by a DFG-grant to MHA (HA 9263/1-1). EHS is moreover grateful for support from CIFAR. The funders had no role in study design, data collection and interpretation, or the decision to submit the work for publication.

## Supplementary Methods

Description of the bioinformatic procedures and tools and Sanger sequencing results.

## Supplementary Tables

Table S1A: List of all crossover and associated gene conversions.

Table S1B: Summary of crossover frequencies per tetrad for the core and accessory chromosomes.

Table S1C: List of genes in crossover hotspots. Table S2A: List of all identified gene conversions.

Table S2B: Summary of the number of SNPs and converted SNPs (total, core (=on core chromosomes), accessory (=on accessory chromosomes)).

Table S2C: List of genes in gene conversion hotspots.

Table S3: Summary of the converted marker and non-converted marker in genomic compartments with the indicated presence/absence of specific histone modifications. Table S4: List of all meiotic *de novo* mutations.

Table S5: Overview of tetrads, spores, mapping results and the number of high-fidelity SNPs.

## Supplementary Figures

**Fig S1.**
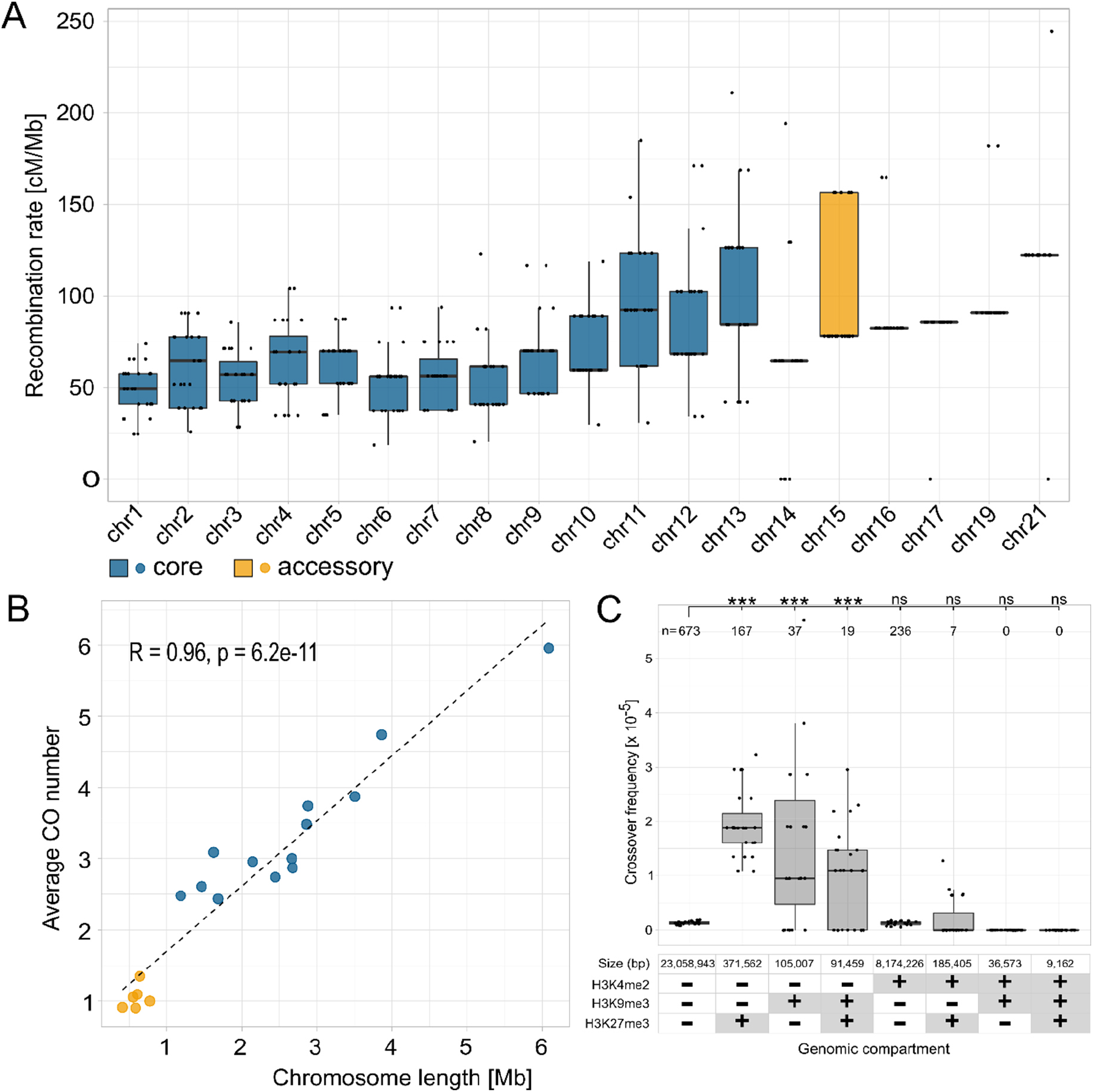
Distribution of recombination rates and the number of crossovers (CO) per chromosome. **A** Recombination rates per chromosome. **B** Average number of CO per chromosome. Box plots in **A** display center line, median; box limits, upper and lower quartiles; whiskers, 1.5x interquartile range; points, rate per tetrad. Points in **B** depict the average CO number for the respective chromosome length; blue points represent core chromosomes; orange points represent accessory chromosomes; the dashed line shows Pearson’s correlation. **C** Correlation between the crossover frequency and chromatin modifications. The presence/absence of the specific chromatin modification (H3K4me2, H3K9me3, or H3K27me3, respectively) in the genomic compartment is depicted with “+”/”-” in the table below the x-axis. The number above each boxplot represents the number of crossovers in the respective genomic compartment. Fisher exact test p-values are shown (*p < 0.05, **p < 0.005, ***p < 0.0005). Box plots display center line, median; box limits, upper and lower quartiles; whiskers, 1.5x interquartile range; points, crossover frequency per genomic compartment per tetrad.

**Fig S2.**
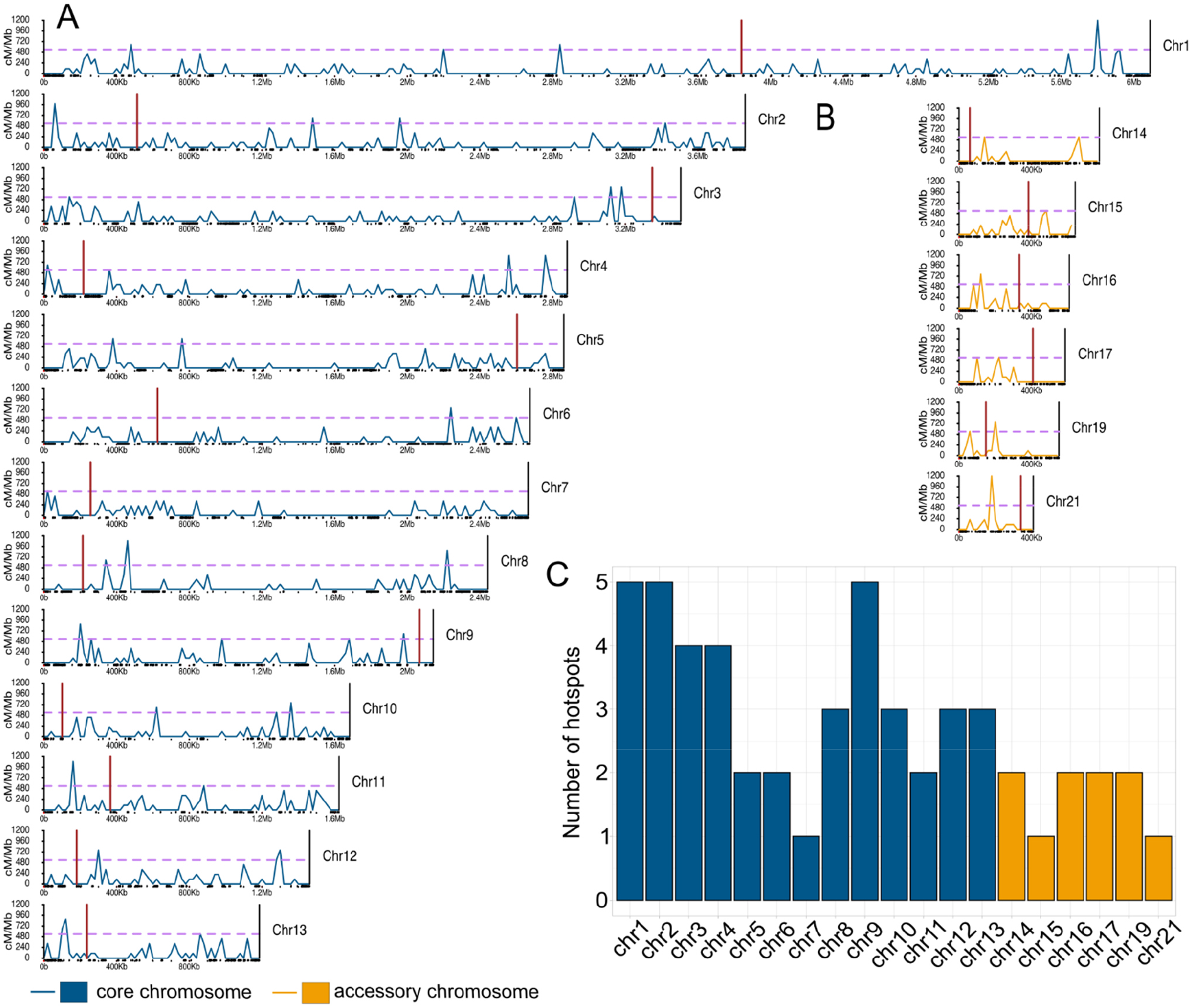
Distribution of recombination rates along the chromosomes in *Z. tritici*. **A** Distribution of recombination rates on core chromosomes. **B** Distribution of recombination rates on accessory chromosomes. Chromosomes are divided in 20 kb non-overlapping windows. The horizontal dashed pink line represent threshold for hotspots with p<0.001 defined by Poisson distribution. The blue lines represent crossover distribution on core chromosomes, orange lines represent crossover distribution on accessory chromosome and red horizontal lines show the centromere position. The x-axis designates the chromosome length. The black rectangles on the x-axis depict transposable elements. **C** Number of hotspots on the indicated chromosomes.

**Fig S3.**
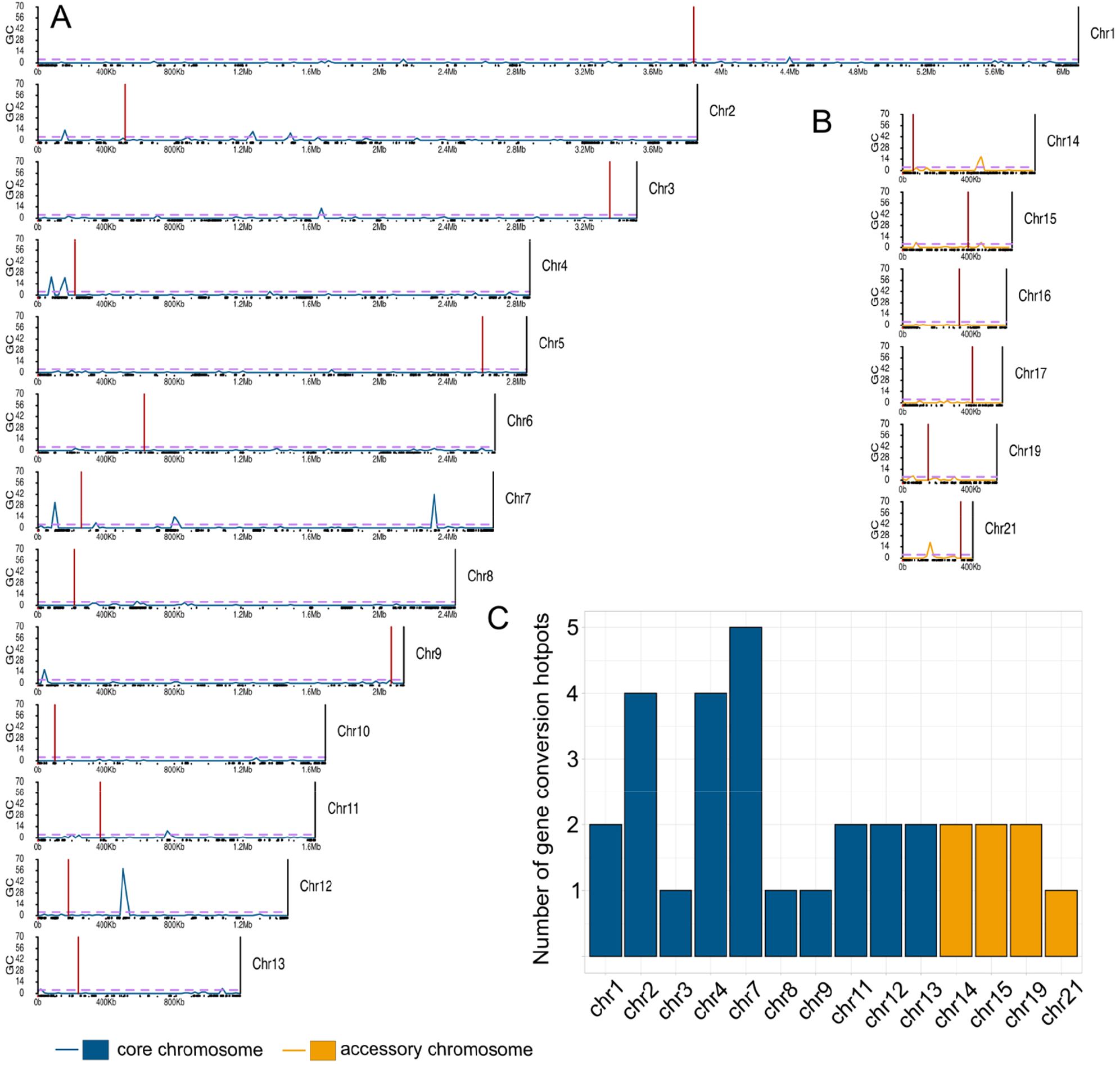
Distribution of gene conversions along the chromosomes in *Z. tritici*. **A** Distribution of gene conversions on core chromosomes. **B** Distribution of gene conversions on accessory chromosomes. Chromosomes are divided in 20 kb non-overlapping windows. The horizontal dashed pink line represent threshold for hotspots with p<0.001 defined by Poisson distribution. The blue lines represent crossover distribution on core chromosomes, orange lines represent crossover distribution on accessory chromosome and red horizontal lines show the centromere position. The x-axis designates the chromosome length. The black rectangles on x-axis depict transposable elements. The y-axis shows the number of gene conversion events (NCO-GC and CO-GC).

**Fig S4.**
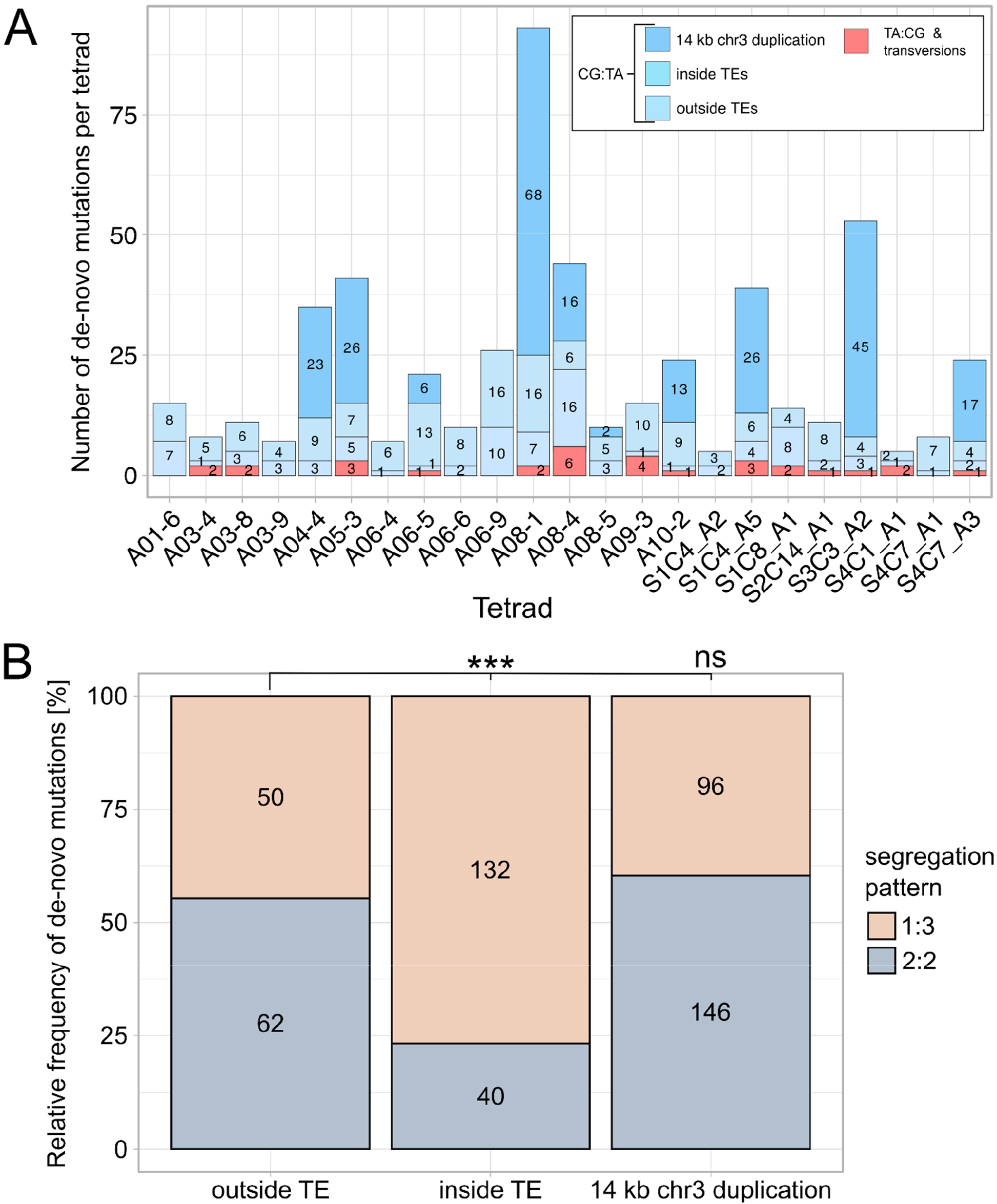
Distribution and segregation of *de novo* mutations. **A** Number of the *de novo* mutations per tetrad. Blue shaded bars display the number of CG:TA transitions on 14 kb duplication on chromosome 3, inside of TEs and outside of TEs per tetrad. Red bars depict the number of TA:CG transitions and transversions. **B** Segregation of *de novo* mutations in different genomic compartments. Light red bars show the relative frequency of *de novo* mutations with 1:3 segregation and grey bars display the relative frequency of *de novo* mutations with 2:2 segregation for the different genomic compartments. Fisher exact test *p-*values are shown (\**p* < 0.05, \*\**p* < 0.005, \*\*\**p* < 0.0005).

## References

1. Peñalba J v., Wolf JBW. 2020. From molecules to populations: appreciating and estimating recombination rate variation. Nat Rev Genet 21:476–492.

2. Keeney S. 2008. Spo11 and the Formation of DNA Double-Strand Breaks in Meiosis, p. 81–123. In Recombination and Meiosis. Springer Berlin Heidelberg, Berlin, Heidelberg.

3. Zelkowski M, Olson MA, Wang M, Pawlowski W. 2019. Diversity and Determinants of Meiotic Recombination Landscapes. Trends in Genetics 35:359–370.

4. Zickler D, Kleckner N. 2015. Recombination, Pairing, and Synapsis of Homologs during Meiosis. Cold Spring Harb Perspect Biol 7:a016626.

5. Chen J-M, Cooper DN, Chuzhanova N, Férec C, Patrinos GP. 2007. Gene conversion: mechanisms, evolution and human disease. Nat Rev Genet 8:762–775.

6. Kleckner N. 1996. Meiosis: how could it work? Proceedings of the National Academy of Sciences 93:8167–8174.

7. Korunes KL, Noor MAF. 2017. Gene conversion and linkage: effects on genome evolution and speciation. Mol Ecol 26:351–364.

8. Youds JL, Boulton SJ. 2011. The choice in meiosis – defining the factors that influence crossover or non-crossover formation. J Cell Sci 124:501–513.

9. Henderson IR, Bomblies K. 2021. Evolution and Plasticity of Genome-Wide Meiotic Recombination Rates. Annu Rev Genet 55:23–43.

10. Bergero R, Ellis P, Haerty W, Larcombe L, Macaulay I, Mehta T, Mogensen M, Murray D, Nash W, Neale MJ, O’Connor R, Ottolini C, Peel N, Ramsey L, Skinner B, Suh A, Summers M, Sun Y, Tidy A, Rahbari R, Rathje C, Immler S. 2021. Meiosis and beyond – understanding the mechanistic and evolutionary processes shaping the germline genome. Biological Reviews 96:822–841.

11. Munz P. 1994. An analysis of interference in the fission yeast Schizosaccharomyces pombe. Genetics 137:701–707.

12. Baudat F, Buard J, Grey C, Fledel-Alon A, Ober C, Przeworski M, Coop G, de Massy B. 2010. PRDM9 is a major determinant of meiotic recombination hotspots in humans and mice. Science 327:836–840.

13. Kauppi L, Jeffreys A, Keeney S. 2004. Where the crossovers are: recombination distributions in mammals. Nat Rev Genet 5:413–424.

14. Haenel Q, Laurentino TG, Roesti M, Berner D. 2018. Meta-analysis of chromosome-scale crossover rate variation in eukaryotes and its significance to evolutionary genomics. Mol Ecol 27:2477–2497.

15. Sardell JM, Cheng C, Dagilis AJ, Ishikawa A, Kitano J, Peichel CL, Kirkpatrick M. 2018. Sex Differences in Recombination in Sticklebacks. G3 (Bethesda) 8:1971–1983.

16. Pan J, Sasaki M, Kniewel R, Murakami H, Blitzblau HG, Tischfield SE, Zhu X, Neale MJ, Jasin M, Socci ND, Hochwagen A, Keeney S. 2011. A Hierarchical Combination of Factors Shapes the Genome-wide Topography of Yeast Meiotic Recombination Initiation. Cell 144:719–731.

17. Borde V, Robine N, Lin W, Bonfils S, Géli V, Nicolas A. 2009. Histone H3 lysine 4 trimethylation marks meiotic recombination initiation sites. EMBO J 28:99–111.

18. Buard J, Barthès P, Grey C, de Massy B. 2009. Distinct histone modifications define initiation and repair of meiotic recombination in the mouse. EMBO J 28:2616–2624.

19. Choi K, Zhao X, Kelly KA, Venn O, Higgins JD, Yelina NE, Hardcastle TJ, Ziolkowski PA, Copenhaver GP, Franklin CH, Mcvean G, Henderson IR. 2013. Arabidopsis meiotic crossover hot spots overlap with H2A.Z nucleosomes at gene promoters. Nature GeNetics VOLUME 45.

20. Underwood CJ, Choi K, Lambing C, Zhao X, Serra H, Borges F, Simorowski J, Ernst E, Jacob Y, Henderson IR, Martienssen RA. 2018. Epigenetic activation of meiotic recombination near Arabidopsis thaliana centromeres via loss of H3K9me2 and non-CG DNA methylation. Genome Res 28:519–531.

21. Stukenbrock EH, Dutheil JY. 2018. Fine-Scale Recombination Maps of Fungal Plant Pathogens Reveal Dynamic Recombination Landscapes and Intragenic Hotspots. Genetics 208:1209–1229.

22. Croll D, Lendenmann MH, Stewart E, McDonald BA. 2015. The Impact of Recombination Hotspots on Genome Evolution of a Fungal Plant Pathogen. Genetics 201:1213–1228.

23. Fouché S, Plissonneau C, McDonald BA, Croll D. 2018. Meiosis leads to pervasive copy-number variation and distorted inheritance of accessory chromosomes of the wheat pathogen Zymoseptoria tritici. Genome Biol Evol 10:1416–1429.

24. Chan AH, Jenkins PA, Song YS. 2012. Genome-Wide Fine-Scale Recombination Rate Variation in Drosophila melanogaster. PLoS Genet 8:e1003090.

25. Daugherty MD, Zanders SE. 2019. Gene conversion generates evolutionary novelty that fuels genetic conflicts. Curr Opin Genet Dev. Elsevier Ltd https://doi.org/10.1016/j.gde.2019.07.011.

26. Lazzaro BP, Clark AG. 2001. Evidence for Recurrent Paralogous Gene Conversion and Exceptional Allelic Divergence in the Attacin Genes of Drosophila melanogaster. Genetics 159:659–671.

27. Thomas JH. Concerted Evolution of Two Novel Protein Families in Caenorhabditis Species https://doi.org/10.1534/genetics.105.052746.

28. Buchmann K. 2014. Evolution of innate immunity: Clues from invertebrates via fish to mammals. Front Immunol 5.

29. Lorenz A, Mpaulo SJ. 2022. Gene conversion: a non-Mendelian process integral to meiotic recombination. Heredity (Edinb) 129:56–63.

30. Mancera E, Bourgon R, Brozzi A, Huber W, Steinmetz LM. 2008. High-resolution mapping of meiotic crossovers and non-crossovers in yeast. Nature 454:479–485.

31. Liu H, Huang J, Sun X, Li J, Hu Y, Yu L, Liti G, Tian D, Hurst LD, Yang S. 2018. Tetrad analysis in plants and fungi finds large differences in gene conversion rates but no GC bias. Nat Ecol Evol 2:164–173.

32. Mansai SP, Kado T, Innan H. 2011. The Rate and Tract Length of Gene Conversion between Duplicated Genes. Genes (Basel) 2:313–331.

33. Lesecque Y, Mouchiroud D, Duret L. 2013. GC-Biased Gene Conversion in Yeast Is Specifically Associated with Crossovers: Molecular Mechanisms and Evolutionary Significance. Mol Biol Evol 30:1409–1419.

34. Marais G. 2003. Biased gene conversion: implications for genome and sex evolution. Trends Genet 19:330–338.

35. Pessia E, Popa A, Mousset S, Rezvoy C, Duret L, Marais GAB. 2012. Evidence for Widespread GC-biased Gene Conversion in Eukaryotes. Genome Biol Evol 4:675–682.

36. Lynch M, Ackerman MS, Gout J-F, Long H, Sung W, Thomas WK, Foster PL. 2016. Genetic drift, selection and the evolution of the mutation rate. Nat Rev Genet 17:704– 714.

37. Narasimhan VM, Rahbari R, Scally A, Wuster A, Mason D, Xue Y, Wright J, Trembath RC, Maher ER, van Heel DA, Auton A, Hurles ME, Tyler-Smith C, Durbin R. 2017. Estimating the human mutation rate from autozygous segments reveals population differences in human mutational processes. Nat Commun 8:303.

38. Rahbari R, Wuster A, Lindsay SJ, Hardwick RJ, Alexandrov LB, al Turki S, Dominiczak A, Morris A, Porteous D, Smith B, Stratton MR, Hurles ME. 2016. Timing, rates and spectra of human germline mutation. Nat Genet 48:126–133.

39. Rattray A, Santoyo G, Shafer B, Strathern JN. 2015. Elevated mutation rate during meiosis in Saccharomyces cerevisiae. PLoS Genet 11:e1004910.

40. Halldorsson B v., Palsson G, Stefansson OA, Jonsson H, Hardarson MT, Eggertsson HP, Gunnarsson B, Oddsson A, Halldorsson GH, Zink F, Gudjonsson SA, Frigge ML, Thorleifsson G, Sigurdsson A, Stacey SN, Sulem P, Masson G, Helgason A, Gudbjartsson DF, Thorsteinsdottir U, Stefansson K. 2019. Human genetics: Characterizing mutagenic effects of recombination through a sequence-level genetic map. Science (1979) 363.

41. Lynch M, Sung W, Morris K, Coffey N, Landry CR, Dopman EB, Dickinson WJ, Okamoto K, Kulkarni S, Hartl DL, Thomas WK. 2008. A genome-wide view of the spectrum of spontaneous mutations in yeast. Proceedings of the National Academy of Sciences 105:9272–9277.

42. Wang L, Sun Y, Sun X, Yu L, Xue L, He Z, Huang J, Tian D, Hurst LD, Yang S. 2020. Repeat-induced point mutation in Neurospora crassa causes the highest known mutation rate and mutational burden of any cellular life. Genome Biol 21:142.

43. Villalba de la Peña M, Summanen PAM, Liukkonen M, Kronholm I. 2022. Variation in spontaneous mutation rate and spectrum across the genome of Neurospora crassa. bioRxiv 2022.03.13.484164.

44. Freitag M, Williams RL, Kothe GO, Selker EU. 2002. A cytosine methyltransferase homologue is essential for repeat-induced point mutation in Neurospora crassa. Proc Natl Acad Sci U S A 99:8802–8807.

45. Galagan JE, Selker EU. 2004. RIP: the evolutionary cost of genome defense. TRENDS in Genetics 20:417–423.

46. Gladyshev E. 2017. Repeat-Induced Point Mutation and Other Genome Defense Mechanisms in Fungi. Microbiol Spectr 5.

47. Gladyshev E, Kleckner N. 2016. Recombination-Independent Recognition of DNA Homology for Repeat-Induced Point Mutation (RIP) Is Modulated by the Underlying Nucleotide Sequence. PLoS Genet 12:e1006015.

48. Selker E. 2002. 15 Repeat-induced gene silencing in fungi, p. 439–450. In Homology Effects. Elsevier.

49. Fudal I, Ross S, Brun H, Besnard A-L, Ermel M, Kuhn M-L, Balesdent M-H, Rouxel T. 2009. Repeat-induced point mutation (RIP) as an alternative mechanism of evolution toward virulence in Leptosphaeria maculans. Molecular Plant-Microbe Interactions 22:932–941.

50. van de Wouw AP, Cozijnsen AJ, Hane JK, Brunner PC, McDonald BA, Oliver RP, Howlett BJ. 2010. Evolution of Linked Avirulence Effectors in Leptosphaeria maculans Is Affected by Genomic Environment and Exposure to Resistance Genes in Host Plants. PLoS Pathog 6:e1001180.

51. Hane JK, Williams AH, Taranto AP, Solomon PS, Oliver RP. 2015. Repeat-Induced Point Mutation: A Fungal-Specific, Endogenous Mutagenesis Process BT – Genetic Transformation Systems in Fungi, Volume 2, p. 55–68. In van den Berg, MA, Maruthachalam, K (eds.),. Springer International Publishing, Cham.

52. Goodwin SB, M’Barek S ben, Dhillon B, Wittenberg AHJ, Crane CF, Hane JK, Foster AJ, van der Lee TAJ, Grimwood J, Aerts A, Antoniw J, Bailey A, Bluhm B, Bowler J, Bristow J, van der Burgt A, Canto-Canché B, Churchill ACL, Conde-Ferràez L, Cools HJ, Coutinho PM, Csukai M, Dehal P, de Wit P, Donzelli B, van de Geest HC, van Ham RCHJ, Hammond-Kosack KE, Henrissat B, Kilian A, Kobayashi AK, Koopmann E, Kourmpetis Y, Kuzniar A, Lindquist E, Lombard V, Maliepaard C, Martins N, Mehrabi R, Nap JPH, Ponomarenko A, Rudd JJ, Salamov A, Schmutz J, Schouten HJ, Shapiro H, Stergiopoulos I, Torriani SFF, Tu H, de Vries RP, Waalwijk C, Ware SB, Wiebenga A, Zwiers LH, Oliver RP, Grigoriev I v., Kema GHJ. 2011. Finished genome of the fungal wheat pathogen Mycosphaerella graminicola reveals dispensome structure, chromosome plasticity, and stealth pathogenesis. PLoS Genet 7.

53. Habig M, Quade J, Stukenbrock EH. 2017. Forward genetics approach reveals host genotype-dependent importance of accessory chromosomes in the fungal wheat pathogen Zymoseptoria tritici. mBio 8.

54. Schotanus K, Soyer JL, Connolly LR, Grandaubert J, Happel P, Smith KM, Freitag M, Stukenbrock EH. 2015. Histone modifications rather than the novel regional centromeres of Zymoseptoria tritici distinguish core and accessory chromosomes. Epigenetics Chromatin 8:41.

55. Habig M, Kema GH, Stukenbrock EH. 2018. Meiotic drive of female-inherited supernumerary chromosomes in a pathogenic fungus https://doi.org/10.7554/eLife.40251.001.

56. Morais D, Gélisse S, Laval V, Sache I, Suffert F. 2016. Inferring the origin of primary inoculum of Zymoseptoria tritici from differential adaptation of resident and immigrant populations to wheat cultivars. Eur J Plant Pathol 145:393–404.

57. Habig M, Lorrain C, Feurtey A, Komluski J, Stukenbrock EH. 2021. Epigenetic modifications affect the rate of spontaneous mutations in a pathogenic fungus. Nat Commun 12:1–13.

58. Möller M, Schotanus K, Soyer JL, Haueisen J, Happ K, Stralucke M, Happel P, Smith KM, Connolly LR, Freitag M, Stukenbrock EH. 2019. Destabilization of chromosome structure by histone H3 lysine 27 methylation. PLoS Genet 15:e1008093.

59. Goodwin SB, M’barek S ben, Dhillon B, Wittenberg AHJ, Crane CF, Hane JK, Foster AJ, van der Lee TAJ, Grimwood J, Aerts A, Antoniw J, Bailey A, Bluhm B, Bowler J, Bristow J, van der Burgt A, Canto-Canche B, Churchill ACL, Conde-Ferraez L, Cools HJ, Coutinho PM, Csukai M, Dehal P, de Wit P, Donzelli B, van de Geest HC, van Ham RCHJ, Hammond-Kosack KE, Henrissat B, Kilian A, Kobayashi AK, Koopmann E, Kourmpetis Y, Kuzniar A, Lindquist E, Lombard V, Maliepaard C, Martins N, Mehrabi R, Nap JPH, Ponomarenko A, Rudd JJ, Salamov A, Schmutz J, Schouten HJ, Shapiro H, Stergiopoulos I, Torriani SFF, Tu H, de Vries RP, Waalwijk C, Ware SB, Wiebenga A, Zwiers L-H, Oliver RP, Grigoriev I v, Kema GHJ. 2011. Finished genome of the fungal wheat pathogen Mycosphaerella graminicola reveals dispensome structure, chromosome plasticity, and stealth pathogenesis. PLoS Genet 7:e1002070.

60. Grandaubert J, Bhattacharyya A, Stukenbrock EH. 2015. RNA-seq-Based Gene Annotation and Comparative Genomics of Four Fungal Grass Pathogens in the Genus Zymoseptoria Identify Novel Orphan Genes and Species-Specific Invasions of Transposable Elements. G3 (Bethesda) 5:1323–1333.

61. Habig M, Kema GHJ, Stukenbrock EH. 2018. Meiotic drive of female-inherited supernumerary chromosomes in a pathogenic fungus. Elife 7.

62. Anderson CM, Chen SY, Dimon MT, Oke A, DeRisi JL, Fung JC. 2011. ReCombine: a suite of programs for detection and analysis of meiotic recombination in whole-genome datasets. PLoS One2011/10/25. 6:e25509–e25509.

63. Cummings WJ, Yabuki M, Ordinario EC, Bednarski DW, Quay S, Maizels N. 2007. Chromatin structure regulates gene conversion. PLoS Biol 5:e246.

64. Bourque G, Burns KH, Gehring M, Gorbunova V, Seluanov A, Hammell M, Imbeault M, Izsvák Z, Levin HL, Macfarlan TS, Mager DL, Feschotte C. 2018. Ten things you should know about transposable elements. Genome Biol 19:199.

65. Wells JN, Feschotte C. 2020. A Field Guide to Eukaryotic Transposable Elements. Annu Rev Genet 54:539–561.

66. Eickbush TH, Malik HS. 2007. Origins and Evolution of Retrotransposons, p. 1111– 1144. In Mobile DNA II.

67. Martin SH, Davey JW, Salazar C, Jiggins CD. 2019. Recombination rate variation shapes barriers to introgression across butterfly genomes. PLoS Biol 17:e2006288.

68. Wang S, Hassold T, Hunt P, White MA, Zickler D, Kleckner N, Zhang L. 2017. Inefficient Crossover Maturation Underlies Elevated Aneuploidy in Human Female Meiosis. Cell 168:977–989.e17.

69. Bhuiyan H, Schmekel K. 2004. Meiotic chromosome synapsis in yeast can occur without spo11-induced DNA double-strand breaks. Genetics 168:775–783.

70. Storlazzi A, Tessé S, Gargano S, James F, Kleckner N, Zickler D. 2003. Meiotic double-strand breaks at the interface of chromosome movement, chromosome remodeling, and reductional division. Genes Dev 17:2675–2687.

71. Tischfield SE, Keeney S. 2012. Scale matters: the spatial correlation of yeast meiotic DNA breaks with histone H3 trimethylation is driven largely by independent colocalization at promoters. Cell Cycle 11:1496–1503.

72. Fowler KR, Sasaki M, Milman N, Keeney S, Smith GR. 2014. Evolutionarily diverse determinants of meiotic DNA break and recombination landscapes across the genome. Genome Res 24:1650–1664.

73. Clouaire T, Legube G. 2015. DNA double strand break repair pathway choice: a chromatin based decision? Nucleus 6:107–113.

74. Jeggo PA, Downs JA. 2014. Roles of chromatin remodellers in DNA double strand break repair. Exp Cell Res 329:69–77.

75. Kalousi A, Soutoglou E. 2016. Nuclear compartmentalization of DNA repair. Curr Opin Genet Dev 37:148–157.

76. Sung P, Klein H. 2006. Mechanism of homologous recombination: mediators and helicases take on regulatory functions. Nat Rev Mol Cell Biol 7:739–750.

77. Alagoz M, Katsuki Y, Ogiwara H, Ogi T, Shibata A, Kakarougkas A, Jeggo P. 2015. SETDB1, HP1 and SUV39 promote repositioning of 53BP1 to extend resection during homologous recombination in G2 cells. Nucleic Acids Res 43:7931–7944.

78. Baldeyron C, Soria G, Roche D, Cook AJL, Almouzni G. 2011. HP1α recruitment to DNA damage by p150CAF-1 promotes homologous recombination repair. Journal of Cell Biology 193:81–95.

79. Lee Y-H, Kuo C-Y, Stark JM, Shih H-M, Ann DK. 2013. HP1 promotes tumor suppressor BRCA1 functions during the DNA damage response. Nucleic Acids Res 41:5784–5798.

80. Soria G, Almouzni G. 2013. Differential contribution of HP1 proteins to DNA end resection and homology-directed repair. Cell Cycle 12:422–429.

81. Sun Y, Jiang X, Xu Y, Ayrapetov MK, Moreau LA, Whetstine JR, Price BD. 2009. Histone H3 methylation links DNA damage detection to activation of the tumour suppressor Tip60. Nat Cell Biol 11:1376–1382.

82. Ayrapetov MK, Gursoy-Yuzugullu O, Xu C, Xu Y, Price BD. 2014. DNA double-strand breaks promote methylation of histone H3 on lysine 9 and transient formation of repressive chromatin. Proc Natl Acad Sci U S A 111:9169–9174.

83. Lemaître C, Grabarz A, Tsouroula K, Andronov L, Furst A, Pankotai T, Heyer V, Rogier M, Attwood KM, Kessler P, Dellaire G, Klaholz B, Reina-San-Martin B, Soutoglou E. 2014. Nuclear position dictates DNA repair pathway choice. Genes Dev 28:2450–2463.

84. Schep R, Brinkman EK, Leemans C, Vergara X, van der Weide RH, Morris B, van Schaik T, Manzo SG, Peric-Hupkes D, van den Berg J, Beijersbergen RL, Medema RH, van Steensel B. 2021. Impact of chromatin context on Cas9-induced DNA double-strand break repair pathway balance. Mol Cell 81:2216-2230.e10.

85. van Wyk S, Wingfield BD, de Vos L, van der Merwe NA, Steenkamp ET. 2021. Genome-Wide Analyses of Repeat-Induced Point Mutations in the Ascomycota. Front Microbiol 11.

86. Coleman JJ, Rounsley SD, Rodriguez-Carres M, Kuo A, Wasmann CC, Grimwood J, Schmutz J, Taga M, White GJ, Zhou S, Schwartz DC, Freitag M, Ma L-J, Danchin EGJ, Henrissat B, Coutinho PM, Nelson DR, Straney D, Napoli CA, Barker BM, Gribskov M, Rep M, Kroken S, Molnar I, Rensing C, Kennell JC, Zamora J, Farman ML, Selker EU, Salamov A, Shapiro H, Pangilinan J, Lindquist E, Lamers C, Grigoriev I v, Geiser DM, Covert SF, Temporini E, Vanetten HD. 2009. The genome of Nectria haematococca: contribution of supernumerary chromosomes to gene expansion. PLoS Genet 5:e1000618.

87. Cuomo CA, Güldener U, Xu J-R, Trail F, Turgeon BG, di Pietro A, Walton JD, Ma L-J, Baker SE, Rep M. 2007. The Fusarium graminearum genome reveals a link between localized polymorphism and pathogen specialization. Science (1979) 317:1400–1402.

88. Graïa F, Lespinet O, Rimbault B, Dequard-Chablat M, Coppin E, Picard M. 2001. Genome quality control: RIP (repeat-induced point mutation) comes to Podospora. Mol Microbiol 40:586–595.

89. Idnurm A, Howlett BJ. 2003. Analysis of loss of pathogenicity mutants reveals that repeat-induced point mutations can occur in the Dothideomycete Leptosphaeria maculans. Fungal Genetics and Biology 39:31–37.

90. Ikeda K, Nakayashiki H, Kataoka T, Tamba H, Hashimoto Y, Tosa Y, Mayama S. 2002. Repeat-induced point mutation (RIP) in Magnaporthe grisea: implications for its sexual cycle in the natural field context. Mol Microbiol 45:1355–1364.

91. Pomraning KR, Connolly LR, Whalen JP, Smith KM, Freitag M. 2013. Repeat-induced point mutation, DNA methylation and heterochromatin in Gibberella zeae (anamorph: Fusarium graminearum). Fusarium Genomics, Molecular and Cellular Biology 93–109.

92. van de Wouw AP, Elliott CE, Popa KM, Idnurm A. 2019. Analysis of Repeat Induced Point (RIP) Mutations in Leptosphaeria maculans Indicates Variability in the RIP Process Between Fungal Species. Genetics 211:89–104.

93. Finnegan DJ. 2012. Retrotransposons. Current Biology 22:R432–R437.

94. Rouxel T, Grandaubert J, Hane JK, Hoede C, van de Wouw AP, Couloux A, Dominguez V, Anthouard V, Bally P, Bourras S, Cozijnsen AJ, Ciuffetti LM, Degrave A, Dilmaghani A, Duret L, Fudal I, Goodwin SB, Gout L, Glaser N, Linglin J, Kema GHJ, Lapalu N, Lawrence CB, May K, Meyer M, Ollivier B, Poulain J, Schoch CL, Simon A, Spatafora JW, Stachowiak A, Turgeon BG, Tyler BM, Vincent D, Weissenbach J, Amselem J, Quesneville H, Oliver RP, Wincker P, Balesdent MH, Howlett BJ. 2011. Effector diversification within compartments of the Leptosphaeria maculans genome affected by Repeat-Induced Point mutations. Nat Commun 2.

95. Frantzeskakis L, di Pietro A, Rep M, Schirawski J, Wu CH, Panstruga R. 2020. Rapid evolution in plant-microbe interactions - a molecular genomics perspective. New Phytol 225:1134–1142.

96. Langmead B, Salzberg SL. 2012. Fast gapped-read alignment with Bowtie 2. Nat Methods 9:357–359.

97. McKenna A, Hanna M, Banks E, Sivachenko A, Cibulskis K, Kernytsky A, Garimella K, Altshuler D, Gabriel S, Daly M, DePristo MA. 2010. The Genome Analysis Toolkit: a MapReduce framework for analyzing next-generation DNA sequencing data. Genome Res 20:1297–1303.

98. Li H, Handsaker B, Wysoker A, Fennell T, Ruan J, Homer N, Marth G, Abecasis G, Durbin R. 2009. The Sequence Alignment/Map format and SAMtools. Bioinformatics 25:2078–2079.

99. Quinlan AR. 2014. BEDTools: The Swiss-Army Tool for Genome Feature Analysis. Curr Protoc Bioinformatics 47.

100. Danecek P, Auton A, Abecasis G, Albers CA, Banks E, DePristo MA, Handsaker RE, Lunter G, Marth GT, Sherry ST, McVean G, Durbin R. 2011. The variant call format and VCFtools. Bioinformatics 27:2156–2158.

101. Layer RM, Chiang C, Quinlan AR, Hall IM. 2014. LUMPY: a probabilistic framework for structural variant discovery. Genome Biol 15:R84.

102. Chiang C, Layer RM, Faust GG, Lindberg MR, Rose DB, Garrison EP, Marth GT, Quinlan AR, Hall IM. 2015. SpeedSeq: ultra-fast personal genome analysis and interpretation. Nat Methods 12:966–968.

